# Distinct synaptic and related transcriptional abnormalities in neonatal, childhood and mature autism model of primate: implications for early-age therapeutic intervention

**DOI:** 10.1101/2020.08.18.255240

**Authors:** Satoshi Watanabe, Tohru Kurotani, Tomofumi Oga, Jun Noguchi, Risa Isoda, Akiko Nakagami, Kazuhisa Sakai, Keiko Nakagaki, Kayo Sumida, Kohei Hoshino, Koichi Saito, Izuru Miyawaki, Masayuki Sekiguchi, Keiji Wada, Takafumi Minamimoto, Noritaka Ichinohe

## Abstract

Autism spectrum disorder (ASD) is a synapse-related disorder that is diagnosed at around 3 years of age. Earlier intervention is desirable for better ASD prognosis; however, there is limited biological literature regarding early-age ASD. This study aimed to assess altered cortical synapses and gene expression in the ASD model marmoset. There were distinct phenotypes in the model animals across the neonate, childhood, and mature stages in the dorsomedial prefrontal cortex (Brodmann area 8b/9). At the neonate stage, synapses were underdeveloped and modulated genes were enriched with synaptogenesis- and ASD-related genes. At the childhood stage, synaptic features and gene expressions associated with experience-dependent circuit remodeling were altered in model animals. At the mature stage, there were synapse overdevelopment and altered gene expression similar to those in human ASD. These early synaptic phenotypes and altered gene expressions could be novel targets of efficient therapy from a young age.

## Introduction

Autism spectrum disorder (ASD) is a highly prevalent neurodevelopmental disorder that affects > 1% of the population. ASD is characterized by impaired social interaction and communication, as well as repetitive behaviors and restricted interests. Currently, its core symptoms cannot be cured and there is a need to develop pharmacological therapy. Patients with ASD typically show delayed development of social and language skills and are diagnosed at around 3 years old. However, early behavioral signs are predictive of pre-existing abnormalities in circuit development. The early years of a child’s life are highly important for subsequent mental health and development. Behavioral intervention, which assists in symptom management of children with ASD, could improve the outcome if began before 4 years of age ^1^. There is a need to identify biological abnormalities underlying behavioral symptoms in order to understand the pathogenesis and develop effective pharmacological treatment starting at early life. However, this has been limited by the lack of human data before the diagnosis age.

ASD is considered a synaptopathy and there is evidence regarding synaptic dysfunctions at both structural and molecular levels. For example, dendritic spines in cortical neurons, which are the postsynaptic structure of excitatory synapses, are affected in patients with ASD ^2,3^. Further, multiple ASD-related genes are involved in synaptic functions ^4^. Synapses connect neurons to form neural circuits, which perform specific brain functions. Normal synaptic development proceeds in a precisely regulated manner, which is indicated by the excessive synaptic formation in the early postnatal stage followed by post-childhood pruning ^5^. Synaptic development involves two processes: genetically programmed synaptogenesis and activity-dependent remodeling. Impairment of these processes could cause abnormal circuit generation and ASD symptoms.

ASD affects primate-specific areas, which is involved in social functions unique to primates (e.g. the dorsolateral frontal cortex) ^6,7^. Further, primates have similar gene expression patterns during development ^8^. Therefore, ASD primate models have advantages over rodent models ^9^. Common marmoset (*Callithrix jacchus*), a small New World monkey, is suitable for studies on neurodevelopmental diseases given the similar cortical architecture and comparable synaptic development to humans ^10,11^. Further, it has the advantages of easy handling, rapid growth, and reproductive efficiency. We have established an ASD marmoset model by administering valproic acid (VPA) *in utero* ^12,13^. VPA, an antiepileptic drug, increases the ASD risk in the offspring ^14^ by inhibiting histone deacetylase (HDAC) and DNA methylation ^15^; further, it has been used to establish ASD model animals ^16,17^. VPA-exposed offspring present with abnormalities in social behavior ^12,13^, axon bundle structure ^18^, and neuroimmune cells ^19^, which are consistent with human data.

From birth to childhood, the brain experiences drastic changes in the neural circuit organization and gene expressions ^5,20^. We hypothesized that ASD affects synapses and gene expressions in an age-specific way during the early postnatal period. We systematically analyzed synaptic structure and activity using acute brain slices from the dorsomedial prefrontal cortex of a VPA-induced ASD model marmoset at the ages of 0, 3, and 6 months (0M, 3M, 6M), which correspond to neonate, childhood, and adolescence, respectively ^10,11^. We observed a clear phenotype transition in synaptic structure and functions throughout this period. Moreover, we examined developmental alterations of the cortical transcriptome in the ASD model marmoset to elucidate the genetic basis of anomalous synaptic development, and found differentially expressed genes (DEGs) that were associated with the specific molecular pathways relevant for ASD and synaptic development. Based on synaptic and molecular analyses, we suggest that there is a unique synaptogenesis impairment at the neonate stage and that circuit remodeling is specifically altered at the childhood stage in patients with ASD. Further, synapses and gene expressions in the mature model marmoset have revealed common dysregulated pathways between the ASD model marmoset and human idiopathic ASD, which accounts for 85% of all ASD cases. Analysis of vocalization at childhood also revealed an impairment characteristic of human ASD. This study suggests the importance of understanding early synaptic phenotypes and the related ASD transcriptome. Further, it suggests novel therapeutic targets for early-age intervention and good outcome throughout life.

## Results

### Early underdevelopment and late overdevelopment of excitatory synapses in VPA-exposed marmosets

Among humans, there is a rapid increase in excitatory synapses and the postsynaptic structure (dendritic spines) in cortical neurons until childhood. This peaks at around 3 years of age and is followed by a decrease through a process called synaptic pruning ^5^. The age of 3 years is the typical diagnosis age of ASD with symptoms becoming evident. Studies on the postmortem brain from patients with ASD have reported a post-childhood increase in the dendritic spine density either due to exaggerated spine formation or incomplete pruning through synaptic development ^2^. Dendritic spines in the prefrontal cortex of typically developed marmosets have shown similar development with a peak spine density at 3M and a subsequent decrease ^10,11^. Therefore, 3M in marmosets could correspond to ∼3 years in humans. To clarify the synaptic phenotypes of VPA-induced ASD model marmosets during synaptogenesis and pruning, we analyzed the excitatory synaptic structure and currents in cortical pyramidal neurons of Brodmann area 8b/9 from VPA-exposed (VPA) and unexposed (UE) marmosets. We focused on layer 3 pyramidal neurons given the relative enrichment of ASD-related gene expression in glutamatergic neurons in the superficial cortical layers in humans ^20^. Accordingly, we made whole-cell recordings from layer 3 pyramidal neurons in acute slices obtained from marmosets prenatally exposed with VPA and UE marmosets at three time points. Further, we morphologically analyzed the neurons after biocytin staining.

The density of dendritic spines on the basal dendrites changed with age. Specifically, it increased from 0M to 3M and then decreased from 3M to 6M in UE animals, which indicates a synaptic production overshoot followed by pruning (Fig. 1a,b). VPA exposure significantly affected spine density development (Fig. 1a,b). Compared to UE animals, the spine density in VPA animals was significantly lower and higher at 0M and 6M, respectively; moreover, it was comparable to that in UE animals at 3M (Fig. 1b; number of data in Table 1). Although the data were not balanced with respect to sex, analysis using the data exclusively from males also provided significant difference between the UE and VPA conditions at 0M (p = 0.0017) and 6M (p = 0.00031), suggesting that the imbalance of the sex ratio is not the cause of the difference. Among VPA animals, there was no significant difference in the spine density at 3M and 6M, which suggests that synaptic pruning is compromised. Contrastingly, there was no significant effect of VPA exposure on dendritic length (Extended Data Fig. 1).

**Fig. 1.**
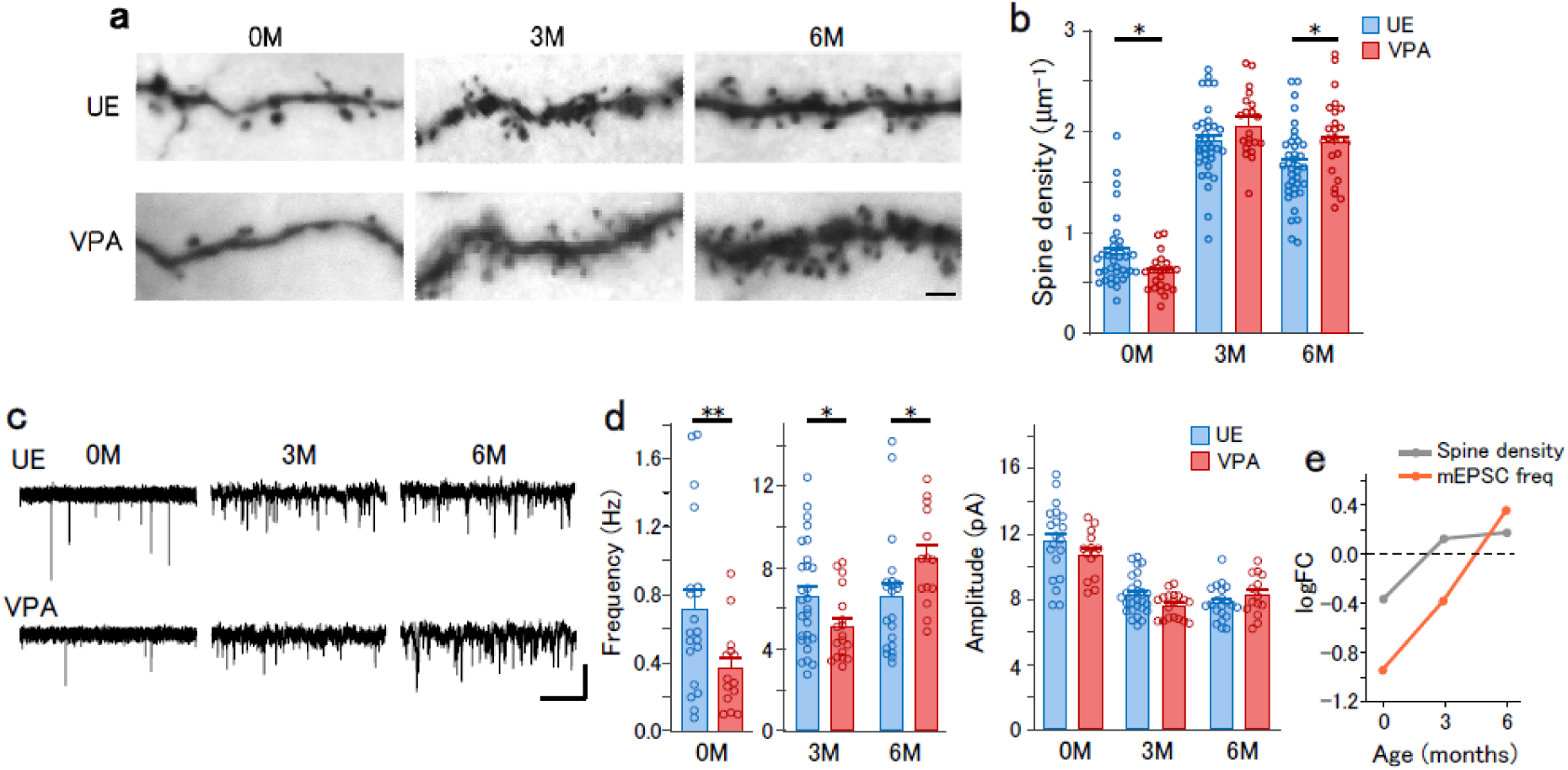
Early underdevelopment and late overdevelopment of excitatory synapses in VPA-exposed marmosets. **a**, Photomicrograph of biocytin-stained dendritic spines in layer 3 pyramidal neurons from UE and VPA animals. Scale bar: 2 µm. **b**, The dendritic spine density in the basal dendrite in layer 3 pyramidal neurons from UE (blue) and VPA (red) animals. Two-way ANOVA, the main effect of age, F (2,184) = 228, p < 2 .2 × 10^−16^; interaction of age × treatment, F (2,184) = 6.94, p = 0.0012. Post-hoc Holm-sidak test in UE animals between 0M and 3M, p = 1.6×10^−21^; between 3M and 6M, p = 0.0053; VPA animals between 3M and 6M, p = 0.081; between UE and VPA animals at 0M, 3M, and 6M, p = 0.015, p = 0.095 and p = 0.019, respectively. *p < 0.05. **c**, Representative traces of mEPSCs recorded from layer 3 pyramidal neurons. Vertical scale bar, 10 pA. Horizontal scale bar, 2.5 s for 0M and 0.5 s for 3M and 6M. **d**, The frequency (left) and amplitude (right) of mEPSCs in UE (blue) and VPA (red) animals. For mEPSC frequency, two-way ANOVA, main effect of age, F (2,102) = 284, p < 2.2 × 10^−16^; interaction of age × treatment, F(2,102) = 9.06, p = 0.00024. Post-hoc Holm-Sidak test in UE animals between 0M and 3M, p = 8.0 × 10^−15^; between 3M and 6M, p = 0.99; between UE and VPA animals at 0M, 3M, and 6M, p = 0.0037, p = 0.038, and p = 0.040, respectively. **p < 0.01, *p < 0.05. For the mEPSC amplitude, main effect of treatment, F(1,102) = 0.26, p = 0.61; interaction of age × treatment, F (2,102) = 1.97, p = 0.15. **e**, logFC values of the spine density and the mEPSC frequency, plotted as a function of age.

**Table 1.** Number of data for each experiment.

|  | UE |  |  |  |  |  | VPA |  |  |  |  |  |  |
| --- | --- | --- | --- | --- | --- | --- | --- | --- | --- | --- | --- | --- | --- |
|  | 0M |  | 3M |  | 6M |  | 0M |  | 3M |  | 6M |  |  |
| Figure | Animals | Samples | Animals | Samples | Animals | Samples | Animals | Samples | Animals | Samples | Animals | Samples | Sample |
| 1b | 5(3/2) | 18/37 | 6(4/2) | 18/37 | 4(2/2) | 23/38 | 3(3/0) | 11/22 | 3(2/1) | 11/21 | 3(2/1) | 9/35 | cells/<br>tissues |
| 1d | 3(2/1) | 20 | 6(3/3) | 28 | 4(1/3) | 21 | 3(2/1) | 14 | 4(3/1) | 18 | 3(2/1) | 14 | cells |
| 2b | 3(2/1) | 21 | 7(4/3) | 37 | 5(2/3) | 25 | 3(2/1) | 16 | 4(3/1) | 25 | 3(2/1) | 16 | cells |
| 2c | 3(2/1) | 20 | 6(3/3) | 26 | 4(1/3) | 17 | 3(2/1) | 14 | 4(3/1) | 16 | 3(2/1) | 11 | cells |
| 2d | 6(4/2) | 19 | 5(4/1) | 22 | 2(1/1) | 11 | 2(2/0) | 9 | 2(1/1) | 15 | 2(1/1) | 11 | cells |
| 3b | 12(7/5) | 22/26 | 5(4/1) | 15/20 | 2(2/0) | 10/16 | 2(2/0) | 11/15 | 5(2/3) | 17/22 | 1(0/1) | 10/15 | slices/<br>tissues |
| 3d,e | 3(2/1) | 10/227 | 5(3/2) | 12/467 | 6(4/2) | 10/472 | 2(2/0) | 12/291 | 2(2/0) | 9/399 | 3(2/1) | 9/406 | cells/<br>tissues |
| 3g |  |  | 2(1/1) | 234 |  |  |  |  | 2(1/1) | 230 |  |  | PSDs |
| 4 |  |  | 8(2/6) |  |  |  |  |  | 5(2/3) |  |  |  |  |
| 5 | 5(2/3) | 10 | 6(4/2) | 9 | 6(5/1) | 9 | 4(1/3) | 11 | 2(1/1) | 6 | 2(1/1) | 6 | tissues |

The numbers in the parentheses for Animals represent the numbers of males/females.

Given that dendritic spines are sites of excitatory synapses, their structural abnormalities in VPA animals could indicate corresponding changes in excitatory synaptic connections. Therefore, we analyzed the miniature excitatory postsynaptic currents (mEPSCs) in acute slices (Fig. 1c). The mEPSCs were blocked by 2,3-dioxo-6-nitro-1,2,3,4-tetrahydrobenzo[f]quinoxaline-7-sulfonamide (NBQX), which confirms that these currents are mediated by AMPA receptors (Extended Data Fig. 2). The mEPSC frequency changed with age (Fig. 1d, left). In UE animals, the mEPSC frequency increased from 0M to 3M; however, it did not significantly differ between 3M and 6M. VPA exposure significantly altered the age-dependent change; specifically, the mEPSC frequency was significantly lower at 0M and higher at 6M in VPA animals. There were no between-group differences in mEPSC amplitude (Fig. 1d, right). These findings demonstrate a transition of VPA-induced alterations in spine density and mEPSCs from underdevelopment at 0M to overdevelopment at 6M, which is summarized in the plots of log fold change (logFC, base-2 logarithm of the ratio of the value in VPA animals to the value in UE animals) (Fig. 1e).

### Reduced E/I ratio in VPA-exposed marmosets at childhood

Excitation-inhibition imbalance is considered an ASD hallmark ^21^. To analyze the E/I ratio, we recorded the miniature inhibitory postsynaptic currents (mIPSCs). The mIPSCs were blocked using picrotoxin, which confirms that these currents are mediated by GABA_A_ receptors (Extended Data Fig. 2). In UE animals, the mIPSC frequency increased with age, which was altered by VPA exposure. Compared with UE animals, VPA animals had a significantly lower and higher mIPSC frequency at 0M and 6M, respectively. However, it was similar to that in UE animals at 3M (Fig. 2a and 2b, left). There was no between-group difference in the mIPSC amplitude (Fig. 2b, right). The ratio of the mESPC to mISPC frequency (E/I ratio of miniature synaptic currents) was affected by VPA exposure (Fig. 2c). Specifically, this ratio was significantly lower in VPA animals at 3M but not at other ages. To determine whether there were alterations in the E/I ratio of stimulus-evoked synaptic transmission, we recorded EPSCs and IPSCs from layer 3 pyramidal neurons in response to field stimulation of layers 4-5 (Extended Data Fig. 3). As expected, VPA animals showed a significantly lower E/I ratio of evoked synaptic currents at 3M but not at other ages (Fig. 2d). The age-dependent modulations of the E/I ratio are shown in the logFC plot (Fig. 2e). Presynaptic properties of excitatory synapses were unlikely to be affected, because VPA exposure did not significantly affect the paired-pulse ratio of EPSCs (Extended Data Fig. 3).

**Fig. 2.**
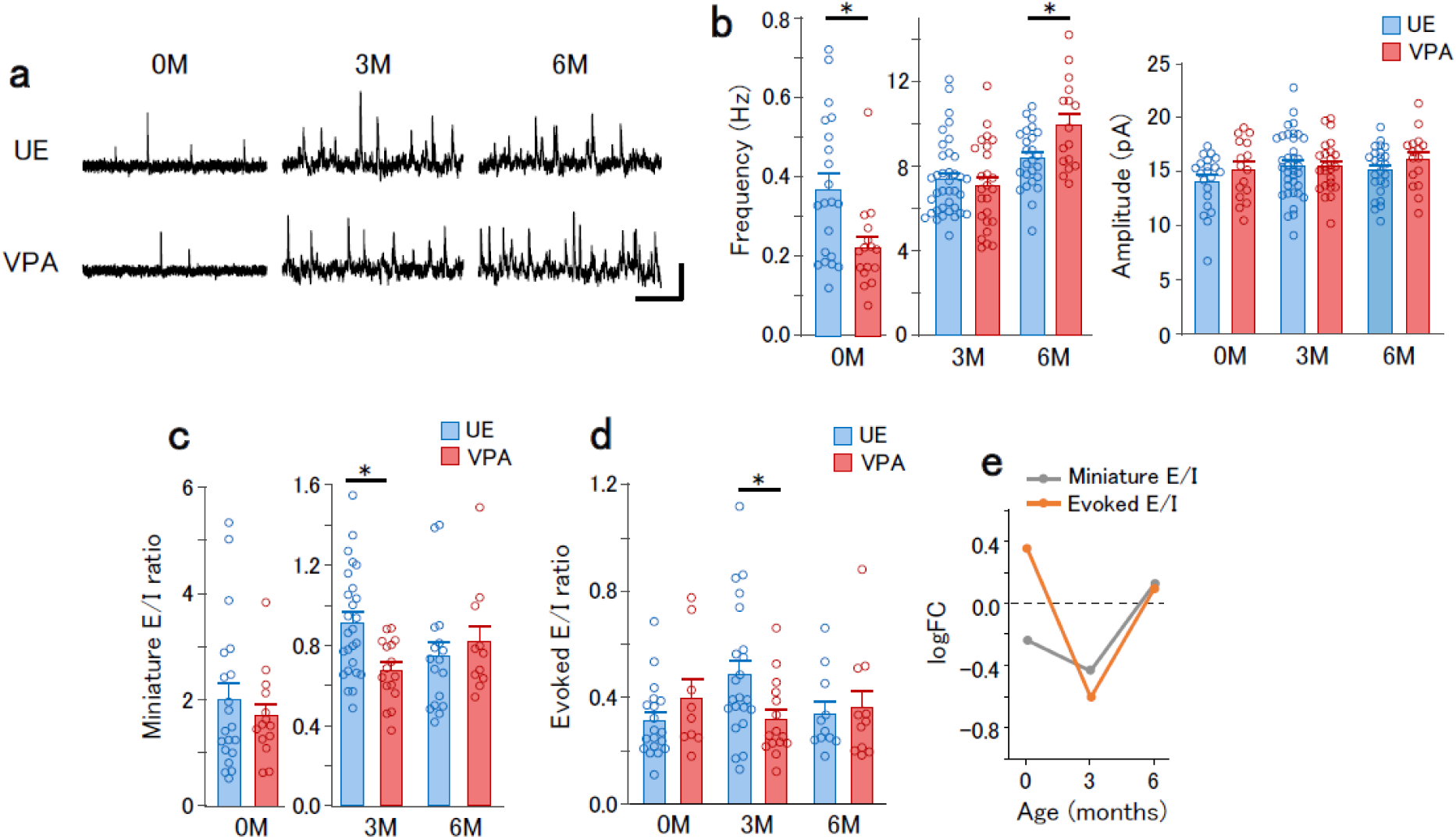
Age-dependent modification of E/I ratio in VPA-exposed marmosets. **a**, Representative mIPSC traces recorded from layer 3 pyramidal neurons. Vertical scale bar, 20 pA. Horizontal scale bar, 2.5 s for 0M and 0.5 s for 3M and 6M. **b**, The frequency (left) and amplitude (right) of mIPSCs in UE (blue) and VPA (red) animals. For mIPSC frequency, main effect of age, F(2,127) = 1216, p < 2.2 × 10^−16^; interaction of age × treatment, F(2,127) = 7.48, p = 0.00085. Post-hoc Holm-Sidak test between UE and VPA animals at 0M, 3M, and 6M, p = 0.011, p = 0.45, and p = 0.027, respectively. *p < 0.05. For mIPSC amplitude, main effect of treatment, F(1,127) = 1.07, p = 0.31; interaction of age × treatment, F(2,127) = 0.42, p = 0.67. **c**, E/I ratio of miniature synaptic currents for UE (blue) and VPA (red) animals. Two-way ANOVA, interaction of age × treatment, F(2,91) = 4.07, p = 0.020. Post-hoc Holm-Sidak test, UE vs. VPA at 0M, 3M, and 6M: p = 0.91, p = 0.018, and p = 0.45, respectively, *p < 0.05. **d**, E/I ratio of the evoked synaptic currents. Two-way ANOVA, interaction of age × treatment, F(2,81) = 3.86, p = 0.025. Post-hoc Holm-Sidak test between UE and VPA at 0M, 3M, and 6M, p = 0.50, p = 0.044, and p = 0.77, respectively, *p < 0.05. **e**, logFC values of the E/I ratio in the frequency of miniature synaptic currents and the evoked synaptic currents, plotted as a function of age.

### Abnormal synaptic plasticity in VPA-exposed marmosets at childhood

The E/I balance in cortical circuits is finely tuned by synaptic plasticity ^22^. The altered E/I ratio in VPA animals observed at 3M suggested downregulation of excitatory synapses at this age; moreover, there might be abnormal regulatory mechanisms of excitatory synapses. Previous studies on ASD model rodents have reported that long-term depression (LTD) is affected ^23^. To determine whether VPA exposure affected LTD of excitatory synapses, we recorded the field EPSPs in layer 3 evoked by layer 4-5 stimulation, and induced LTD by low frequency stimulation (LFS, 1 Hz, 15 min). VPA exposure affected LFS-induced LTD in an age-dependent manner. At 0M, LFS induced similar LTD magnitudes in UE and VPA animals. At 3M, LFS induced LTD in VPA animals, but not UE animals. At 6M, LFS did not induce LTD in either UE or VPA animals (Fig. 3a,b). The logFC value for the field EPSP amplitudes after LFS showed a reduction at 3M (Fig. 3c). These results suggest that synapse susceptibility to LFS is transiently altered at 3M.

**Fig. 3.**
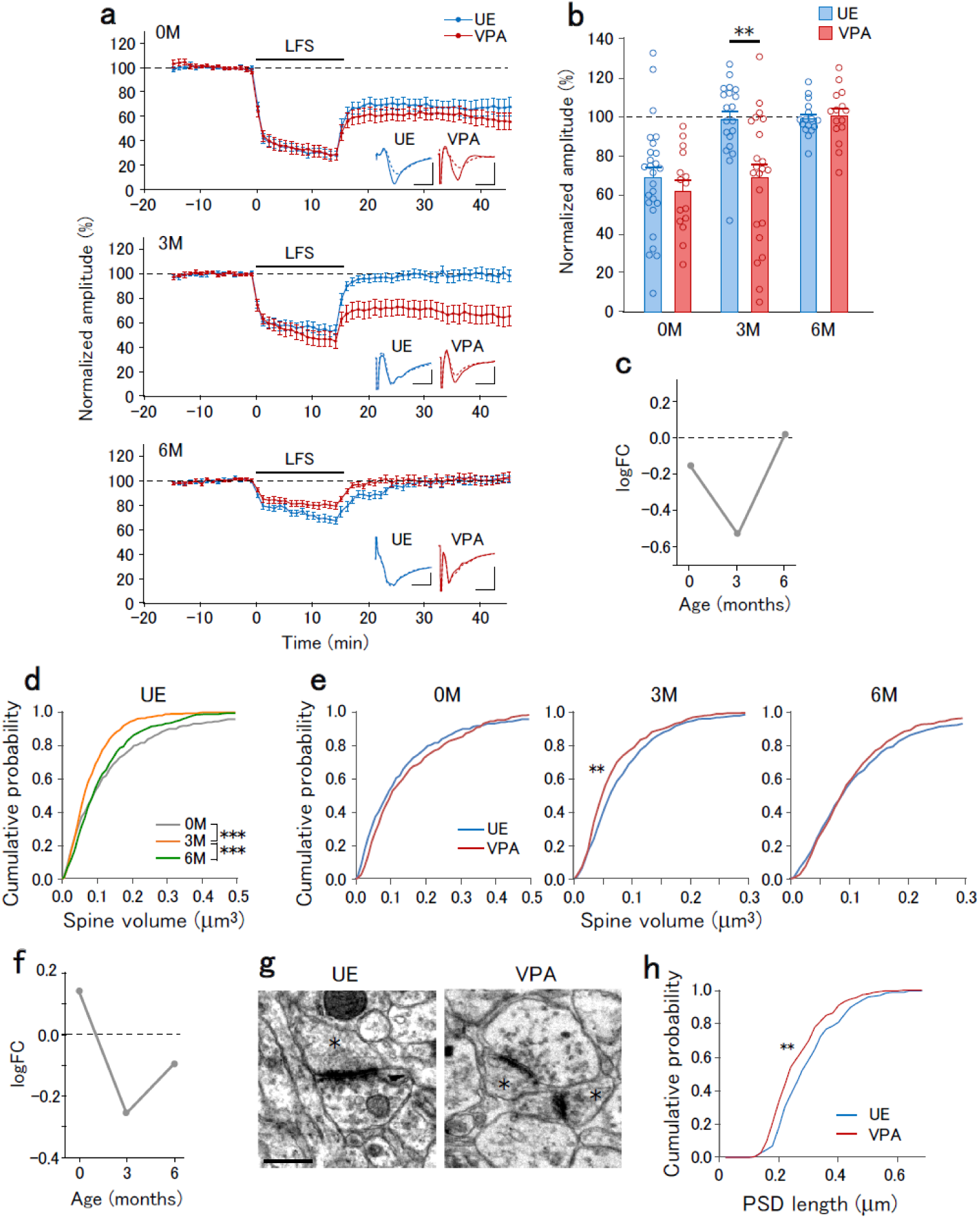
Age-dependent modification of LTD and spine volume in VPA-exposed marmosets. **a**, The time course of normalized field EPSP amplitudes before and after LFS onset that were recorded from slices obtained at 0M (top), 3M (middle), and 6M (bottom) of UE (blue) and VPA (red) animals. Insets represent the average traces of field EPSPs before (−10 to 0 min, solid lines) and after (30 to 40 min, dotted lines) LFS. **b**, Normalized field EPSP amplitudes in UE and VPA animals at post-LFS 30-40 min. Two-way ANOVA, main effect of treatment, F(1,79) = 7.03, p = 0.0092; interaction of age × treatment, F(1,79) = 4.56, p = 0.013. The post-hoc Holm-Sidak test between UE and VPA animals at 0M, 3M, and 6M, p = 0.43, p = 0.0022, and 0.66, respectively. **p < 0.01. **c**, logFC values for the post-LFS normalized field EPSP amplitudes plotted as a function of age. **d**, Cumulative distribution of spine volume in UE animals at 0M, 3M, and 6M. The Kolmogorov-Smirnov test; 0M vs. 3M, p = 6.0 × 10^−5^; 3M vs. 6M, p = 4.7 × 10^−5^; 0M vs. 6M, 0.28. ***p < 0.001. **e**, Cumulative distribution of spine volume in UE (blue) and VPA (red) animals at 0M, 3M, and 6M. The Kolmogorov-Smirnov test between UE and VPA animals at 0M, 3M, and 6M, p = 0.089, 0.0025, and 0.64, respectively, **p < 0.01. **f**, logFC values for the spine volume plotted as a function of age. **g**, Representative electron micrograph of synapses in UE and VPA animals at 3M. Asterisks indicate dendritic spines. Scale bar, 0.5 µm. **h**, Cumulative distribution of the PSD length in UE and VPA animals at 3M. Kolmogorov-Smirnov test, **p = 0.0026.

LFS-induced LTD could be dependent on NMDA receptors given that it was blocked by D-2-amino-5-phosphonovaleric acid (D-APV) (Extended Data Fig. 4). Metabotropic glutamate receptor (mGluR)-dependent LTD induced by (S)-3,5-dihydroxyphenylglycine (DHPG), which is a group I mGluR agonist that induces LTD in the visual cortex ^24^, was observed in both UE and VPA animals at 0M but not at 3M (Extended Data Fig. 4). Therefore, prenatal VPA exposure does not affect mGluR-dependent LTD.

LTD causes dendritic spine shrinkage ^25^, and therefore we analyzed the dendritic spine volume (Extended Data Fig. 5). Both UE and VPA animals displayed an age-dependent alteration in the spine volume distribution (Fig. 3d). Compared with UE animals, VPA animals showed significantly lower spine volume distribution at 3M, but not at 0M and 6M (Fig. 3e). The logFC value for the average spine volume was reduced at 3M (Fig. 3f). Electron microscopy confirmed the VPA exposure-induced synaptic structure impairment at 3M (Fig. 3g). Compared with UE animals, VPA animals showed a significantly lower length of postsynaptic density (PSD) (Fig. 3h).

In summary, prenatal VPA exposure affects LTD, E/I ratio, and spine volume at 3M with the effects at other ages being weaker. This suggests that 3M is another crucial time point in ASD development.

### Altered vocalization in VPA-exposed marmosets at childhood

Abnormal language development is characteristic of ASD; moreover, the dorsomedial prefrontal cortex is among the brain regions involved in vocalization in primates ^26^. A marmoset is a highly social animal and communicates with other marmosets using vocalization. Vocal development in marmosets requires parental vocal feedback ^27^. We postulated that impaired early synaptic development could affect vocal maturation at 3M in marmosets, which corresponds to the childhood stage when language deficits become evident in humans.

We recorded the calls of marmosets in an isolated environment for 5 min and annotated each call (Fig. 4a). The total call number during the isolation tended to be lower in VPA animals; however, there was no significant difference (Fig. 4b). UE marmosets produced various calls with phee calls being the most frequent and constituting 44% of all calls. VPA animals produced a higher ratio of phee call (79%), with lower frequencies for other call types (Fig. 4c). The predominance of the phee call in VPA animals caused a significantly lower entropy of the call types in VPA animals than in UE animals (Fig. 4d). These findings are indicative of the existence of behavioral abnormality in VPA animals following abnormal synaptic development between birth and childhood.

**Fig. 4.**
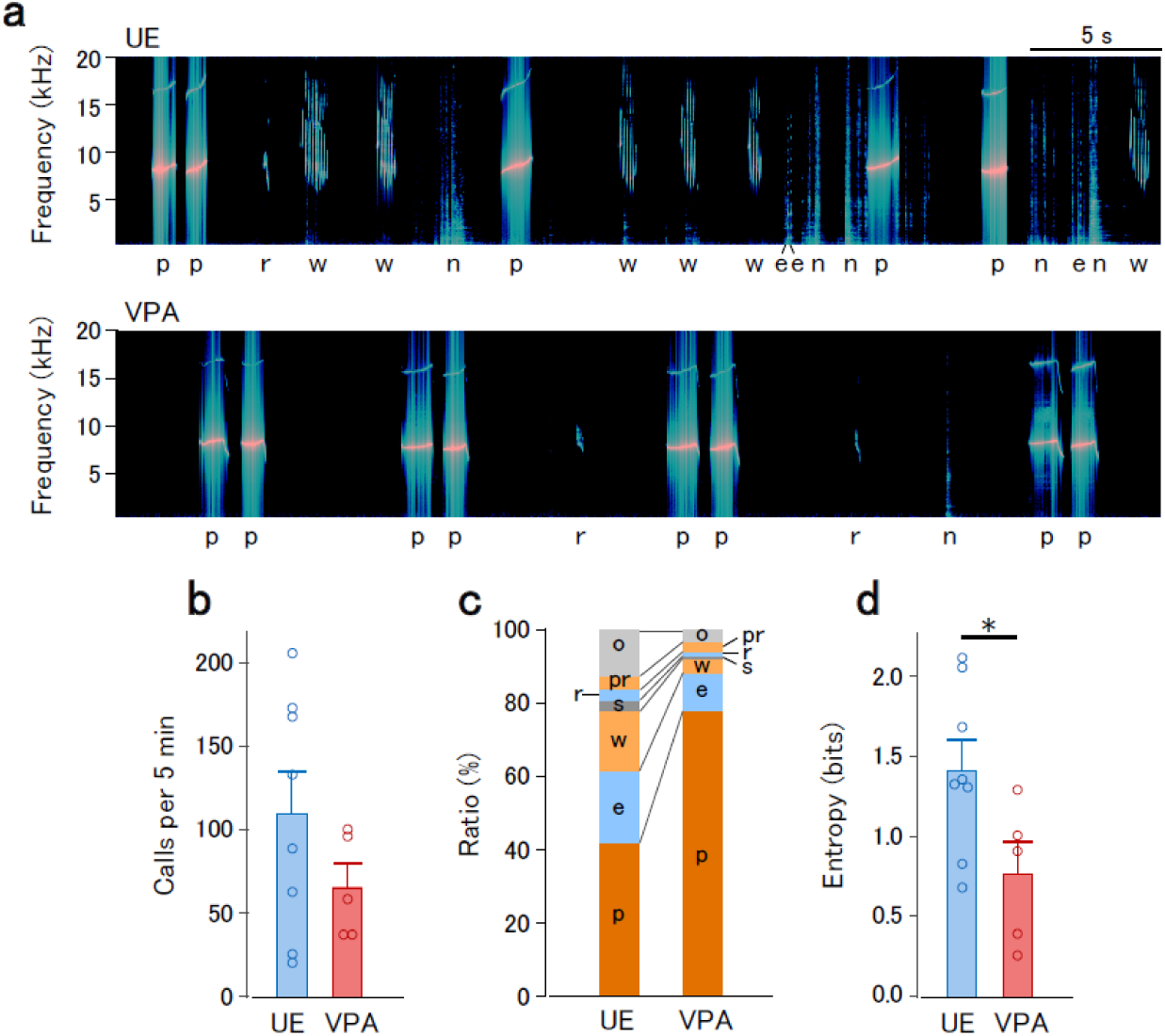
Altered isolation calls in VPA-exposed marmosets. **a**, Representative vocalization spectrogram for UE (top) and VPA (top) animals at 12 weeks of age. The type of call is shown below the spectrogram. Call types: e, ekk or cough; p, phee; r, trill; w, twitter; n indicates noise. **b**, The average number of calls in UE and VPA animals at 11-13 weeks. *t*-test, p = 0.16. **c**, Average ratio of call types in UE and VPA animals. Call types: pr, phee-trill or trill-phee; s, tsik; o, others. Other annotations are as in (a). **d**, The average entropy of calls in UE and VPA animals. *t*-test, *p = 0.036.

### Altered gene expressions in VPA-exposed marmosets

We observed excitatory synapse underdevelopment at 0M, which became overdeveloped at 6M. Further, at 3M, there were reduced E/I ratio and enhanced LTD. We speculated that these phenotypes are accompanied by altered gene expressions. We searched for VPA-modulated gene expression using a custom-made microarray. Among 7951 protein-coding genes expressed in the three cortical regions (areas 8, 12 and TE), there were 1032 DEGs with an absolute value of logFC of > 0.4 and a p_adj_ of < 0.05 at either age.

Since there were synaptic phenotype alterations with age, we analyzed the temporal profile of gene expression alterations. The temporal profile of logFC values revealed distinct VPA-induced modulations. Using the *k*-means clustering method, the DEGs were grouped into 3 clusters as indicated by the colored dots (Fig. 5a). These clusters were discernibly separated on the plot of logFC values at 3M against those at 0M. We have provided a comprehensive list of genes in each cluster (Supplementary Table 1).

**Fig. 5.**
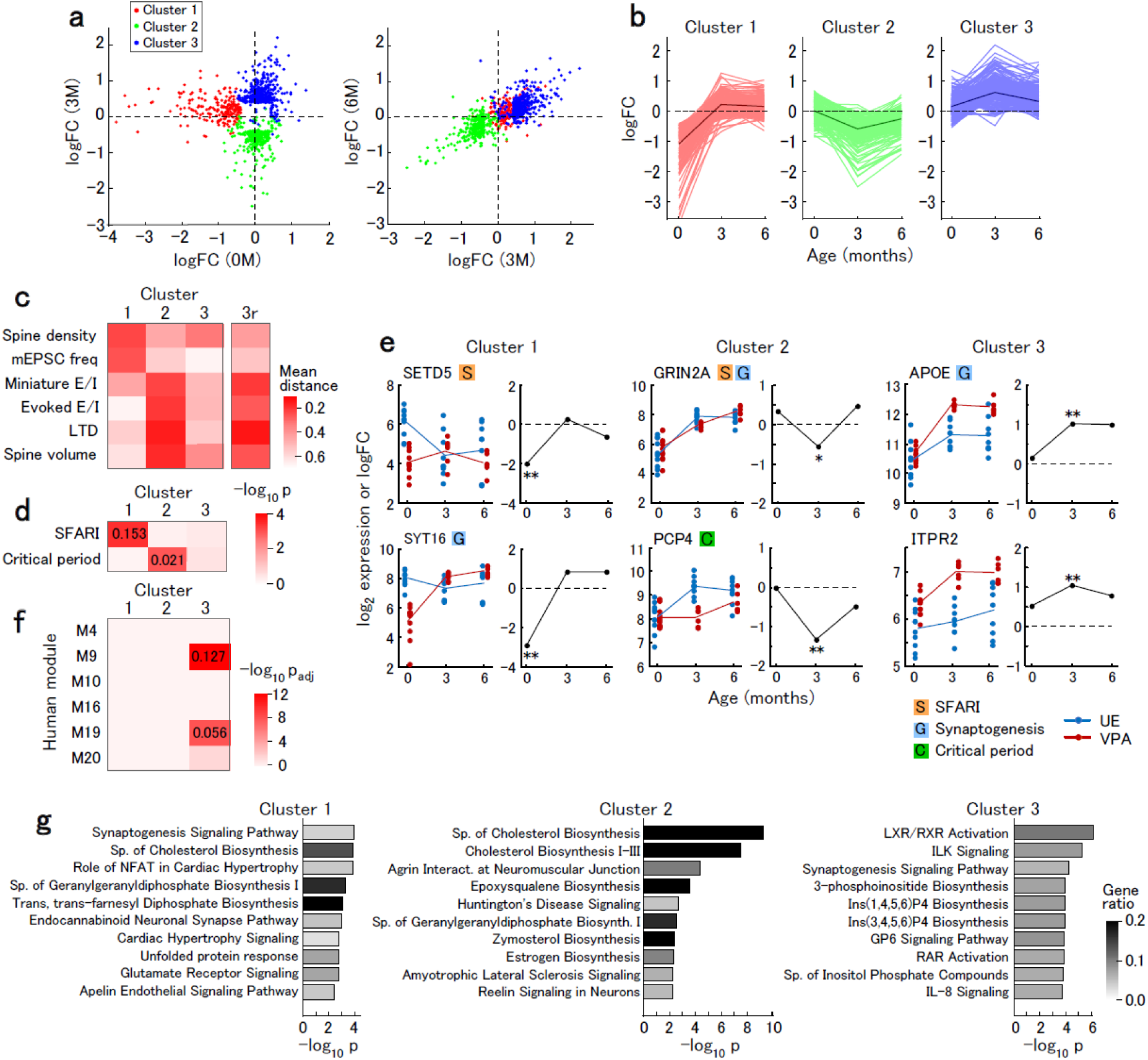
Microarray analysis of differential gene expressions. **a**, Distribution of the logFC values of gene expressions significantly modulated by VPA exposure. The logFC value at 3M was plotted against the logFC value at 0M (left) while the logFC value at 6M was plotted against the logFC value at 3M (right). Each gene was colored based on the cluster. **b**, The time course of logFC values. The traces in light colors represent data from individual genes in the cluster. The traces in dark colors represent the average of all genes in the cluster. **c**, The average distance between the average logFC values for each cluster and the logFC vlues of structural and physiological parameters (spine density, mEPSC frequency, E/I ratio of miniature and evoked synaptic currents, post-LTD field EPSP amplitudes, and spine volume). The column labeled “3r” shows the average distance between the negative average logFC values of cluster 3 and the structural and physiological parameters in order to highlight parameters inversely correlated with cluster 3. **d**, Enrichment of SFARI and critical period-related genes in each cluster. Colors indicate the p values using Fisher’s exact test, while the numbers represent the ratio of SFARI or critical period-related genes in each cluster. Regarding SFARI genes, the number of genes and p value were 31 genes (p = 0.00066), 25 genes (p = 0.25), and 35 genes (p = 0.79), respectively. Regarding critical period-related genes, 0 genes (p = 0.64), 8 genes (p = 0.0026), and 3 genes (p = 1.0), respectively. **e**, The time courses of the log_2_ expressions in UE (blue) and VPA (red) animals (left) and logFC values (right) of representative genes in each cluster. SFARI (S), synaptogenesis-related (G), or critical period-related (C) genes are marked. Asterisks indicate significant differences in gene expressions between UE and VPA animals (**p_adj_ < 0.01, *p_adj_ < 0.05). **f**, Enrichment of genes belonging to each cluster in human ASD-related modules ^31^. Colors indicate adjusted p values using Fisher’s exact test, while the numbers indicate the ratio of the cluster genes that fall in each module. **g**, Enriched pathway terms for each cluster. The color of the bars represents the ratio of the cluster genes in the pathway. Abbreviations: Sp, superpathway; Ins(1,4,5,6)P4, D-myo-inositol (1,4,5,6)-tetrakisphosphate; Ins(3,4,5,6)P4, D-myo-inositol (3,4,5,6)-tetrakisphosphate.

All logFC values of cluster 1 genes were substantially low at 0M (Fig. 5b). The logFC values of cluster 2 and 3 genes were negative and positive at 3M, respectively, while those at 0M and 6M were smaller in magnitude (Fig. 5b). The time course of the logFC values for cluster 1 genes corresponded to those of the spine density and mEPSC frequency, as indicated by the low mean distance between the plots of logFC values of the synaptic parameters and gene expressions (Fig. 5c). This suggests that cluster 1 genes are associated with abnormal synaptic development. The time course of the logFC values for cluster 2 genes was consistent with those of E/I ratios, the post-LTD field EPSP amplitudes, and spine volume. Although the time course of the logFC values for cluster 3 genes were not correlated with the synaptic modulations, the plot of the negative of the logFC values of cluster 3 genes was parallel to the logFC of E/I ratios, post-LTD field EPSP amplitudes, and spine volume (“3r” in Fig. 5c). These results suggest that the expression of the genes in clusters 2 and 3 are associated with synaptic modulations that specifically occur at 3M (Fig. 5c).

Cluster 1, but not clusters 2 and 3, was significantly enriched with ASD-associated (SFARI) genes (Fig. 5d). Further, we analyzed critical period-related gene expressions, since critical period plasticity is affected in ASD model animals ^28^. Cluster 2, but not clusters 1 and 3, was enriched with critical period-related genes with the highest expression at the critical period in the visual cortex of mice ^29,30^ (Fig. 5d, Supplementary Table 2). These findings suggest that genes affected by VPA exposure at different times were probably involved in different aspects of synaptic modulations in patients with ASD. Fig. 5e has several gene expression profiles. Comparison of the gene clusters with modules from weighted gene coexpression network analysis (WGCNA) in human ASD and control samples ^31^ revealed that cluster 3 genes were enriched in two ASD-related modules (Fig. 5f; Supplementary Table1). Module 9 and 19 are associated with astrocytes and microglia; moreover, similar to the marmoset cluster 3 genes, genes in both modules are upregulated in human ASD. Astrocytes and microglia are cortical components that regulate synapses. Pathway analysis revealed that the synaptogenesis signaling pathway was the 1st, 38th, and 3rd pathway term of cluster 1, 2, and 3, respectively, with the enrichment of this pathway being significant in these clusters. The superpathway of cholesterol biosynthesis was the 3rd and 1st pathway term of cluster 1 and 2, respectively (Fig. 5g, Supplementary Table 3).

Finally, we compared gene expression modulation in VPA-exposed marmosets with that in human ASD (Fig. 6a). There was a significant correlation of the logFC values of modulated genes at 3M in marmosets with those in human ASD ^31^. Further, there was a correlation of the logFC values at 6M with human genes; however, fewer genes were modulated at this age. Contrastingly, there was no correlation of logFC at 0M with that of human ASD (Fig. 6a). These results suggest that modulated gene expressions in marmosets after 3M are consistent with those observed in human idiopathic ASD. Contrastingly, there is a distinct pattern of modulated gene expressions in neonatal marmosets, which could be attributed to the fact that available human ASD data have been obtained from juvenile to adult samples. Gene expression modulations in neurons derived from induced pluripotent stem cells (iPSCs) from patients with ASD, which correspond to mid-fetal stage, should differ from those observed in postmortem samples ^32^. Indeed, we observed a negative correlation between the gene expression modulations in marmoset at 3M and those in iPSC-derived neurons from humans ASD ^33^ (Fig. 6b). In comparison, the correlation between marmoset at 0M and iPSC-derived neurons was lower (Fig. 6b). These results suggest that gene expression modulations at the perinatal stage are different from those at both fetal and mature ages. There were higher correlations in the logFC values between the VPA-exposed marmosets at 3M or 6M and human ASD compared to those reported between VPA-exposed rats ^34^ and human ASD (Fig. 6c). This suggests that the marmoset VPA model mimics human ASD at the molecular level more closely than the rat VPA model.

**Fig. 6.**
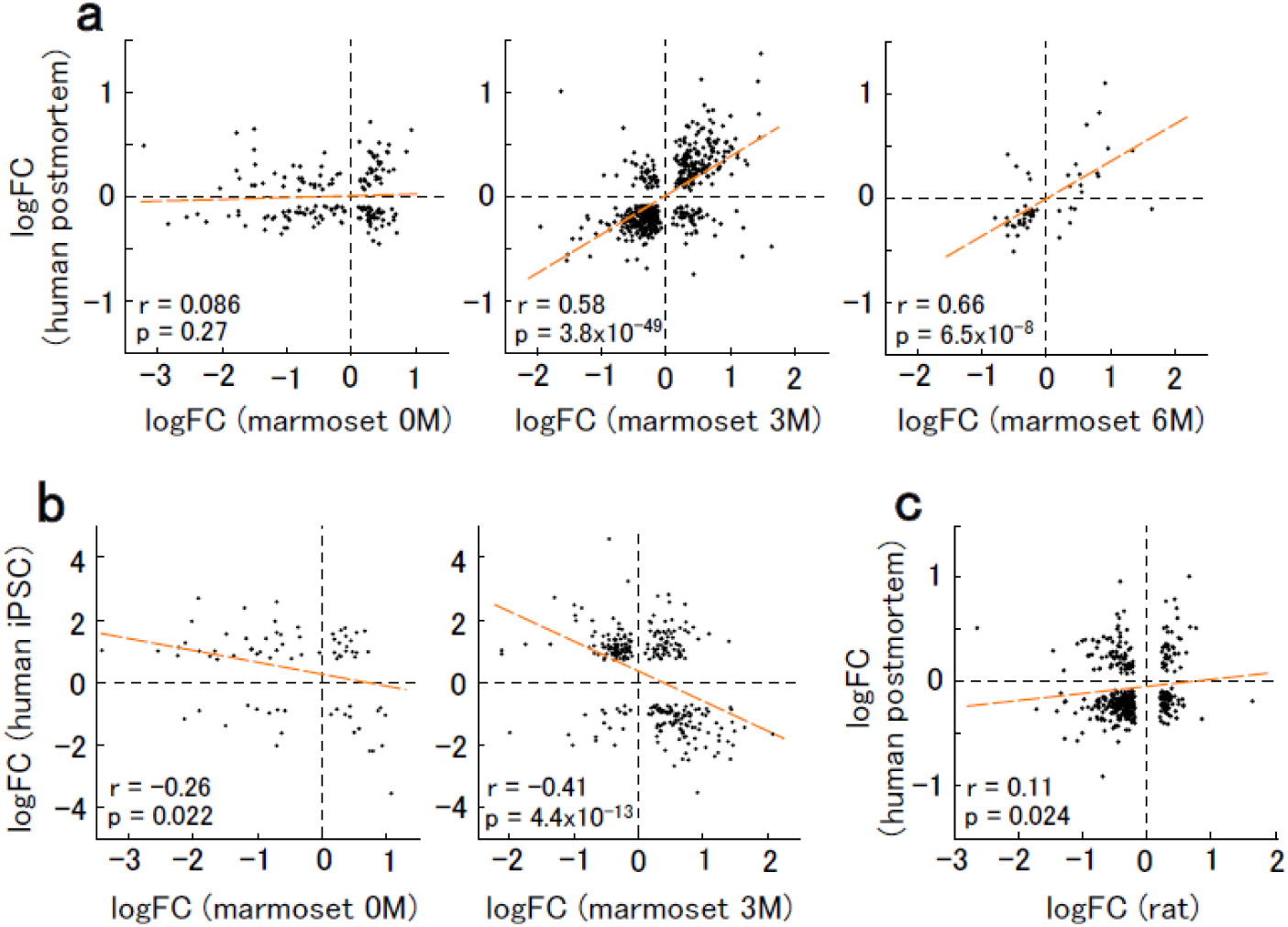
Relationship between gene expression modulations in VPA-exposed marmosets and human ASD. **a**, Relationship between the logFC values in marmoset and postmortem human ASD ^31^. Modulated genes with p_adj_ < 0.1 are plotted. For each plot, Spearman’s correlation coefficients (r) and p-values (p) are shown. Dashed orange line is the linear regression. **b**, Relationship between the logFC values in marmosets and iPSC-derived neurons from human ASD (31 days of terminal differentiation) ^33^. Modulated genes with p_adj_ < 0.1 are plotted. **c**, Relationship between the logFC values in VPA-exposed rats and postmortem human ASD ^34^. Modulated genes with p_adj_ < 0.05 are plotted.

## Discussion

This study aimed to determine abnormalities in synaptic features and related transcriptome from birth to childhood, which is when ASD symptoms become diagnosable and a foundation for future mental health is established. Consequently, we analyzed the synaptic phenotypes and related-transcriptome in VPA-exposed ASD model marmosets at the neonate, childhood, and adolescence. Our results suggest the existence of distinct synapse pathophysiology and gene expressions among different developmental stages. At the neonate stage, there were reduced spine density and mEPSC frequency in the ASD model marmoset, which suggests reduced synaptogenesis in early postnatal development. Moreover, transcriptome analysis revealed that one DEG cluster (cluster 1) was specifically downregulated at the neonate stage. These genes were associated with synaptogenesis signaling and enriched with SFARI genes, which further indicates that synaptogenesis is compromised at the neonate stage. At the childhood stage, there was a reduced E/I ratio and enhanced LTD of excitatory synapses in the ASD model marmoset. LTD during childhood is an essential component of circuit remodeling. Transcriptome analysis revealed two DEG clusters that were specifically modulated at childhood. Cluster 2, which was downregulated at childhood, was enriched with critical period-related genes, suggesting that circuit remodeling may be affected during childhood. Synaptic overdevelopment in mature model marmoset was similar to that in human ASD. Moreover, there was a good correlation of post-childhood gene expression alterations with human ASD. This suggests that phenotypes in mature model marmosets are consistent with observations in post-mortem brains obtained from mostly adult patients with idiopathic ASD. Synaptic phenotypes and gene expression modifications at the neonate and childhood stage were consistent with the findings observed in neurons derived from iPSCs. These results further validate common disordered pathways between the ASD model marmoset and individuals with idiopathic ASD.

Previous studies have reported abnormal dendritic spine development in human ASD ^2,3^. Although the spine density is significantly higher in patients with ASD at 13-19 years of age (adolescence), there was no significant difference observed at the age of 2-9 years (childhood). This indicates that the effect on spine density worsens with age as synaptic pruning is compromised ^3^. Consistently, we observed an elevated spine density in VPA-exposed marmosets at 6M (adolescence). Contrastingly, the spine density was lower at 0M. There was reduced synaptogenesis in iPSC-derived neurons from patients with ASD, which corresponded to the mid-fetal time ^35^. This suggests that early synaptic maldevelopment and late overdevelopment are common phenotypes to VPA-exposed marmosets and human ASD.

ASD is thought to involve a mechanism similar to the critical period plasticity ^28^. LTD is required for ocular dominance plasticity in the visual cortex ^36^, which suggests that the effect of VPA exposure on LTD, specifically at childhood, could affect circuit remodeling. Structure of dendritic spines is critical for AMPA receptor expression ^37^. Small spines and filopodia, which were abundant in VPA animals at 3M, may be the structural substrates of silent synapses. These contribute to the reduced E/I ratio among childhood regardless of unaltered spine density (Fig. 1). The E/I ratio is a common circuit property that is affected in ASD animal models ^21^. The E/I ratio is an essential factor that regulates the critical period plasticity; further, it is involved in homeostatic synapse regulation in circuit remodeling ^22^. Reduced synaptogenesis in neonates and E/I ratio at childhood may restrict the transmission of social and communication information required for appropriate experience-dependent circuit development as observed in sensory deprivation ^38^. This affects the developmental trajectory and causes circuit malfunctioning that subsequently causes ASD.

Language acquisition becomes progressively difficult after a few years of age, which suggests that early experience during the sensitive period is necessary for language development ^39^. Most patients with ASD have difficulty in context-dependent communication and often produce stereotyped repetitive speech during childhood. There is an overlap of ASD symptom emergence with the sensitive periods of social and language skills. Moreover, LTD is required for vocal learning in songbirds ^40^. Consistently with compromised synaptic development and abnormal LTD, VPA animals showed abnormal vocalization at childhood (Fig. 4). The increased phee call ratio and decreased vocalization variability in VPA-exposed marmosets at childhood is similar to human ASD and suggests a close relationship between vocalization and synaptic phenotypes until this age.

Transcriptomic analysis revealed altered gene expressions in VPA animals. We classified the affected genes into 3 clusters based on the time courses of logFC values. Although clustering was performed independently of the synaptic data, each cluster was closely related with synaptic phenotypes (Fig. 5). This suggests that these clusters represent the molecular mechanisms underlying early pathological ASD development. Cluster 1 genes were downregulated in neonates; additionally, the temporal modulation pattern was similar to that of the spine density and the frequency of miniature synaptic currents. This suggests an association of altered gene expressions with abnormal synaptic development. Indeed, the synaptogenesis signaling pathway was the top pathway term for this cluster. Cluster 1 was enriched with SFARI genes. This cluster contained SETD5 (SET Domain Containing 5), which is a SFARI gene involved in DNA methylation and suggests a common epigenetic mechanism between the VPA model marmosets and ASD humans ^41^. Clusters 2 and 3 genes were downregulated and upregulated at 3M, and these temporal modulation patterns were similar and inverse to alterations in the E/I ratio, the post-LTD field EPSP amplitude, and spine volume. GRIN2A (NMDA receptor subunit 2A), which was among the cluster 2 genes, has been suggested to be associated with ASD and other neurological diseases ^42^. Moreover, it may be a key molecule for the altered synaptic properties in VPA animals. GRIN2A expression is downregulated by reduced sensory stimuli ^43^, and GRIN2A-knockout mice show an enhanced LTD with weak stimuli ^44^. Further, cluster 2 included critical period-related genes showing the highest expression in the critical period (about 4 weeks of age) in rodents ^29,30^, including PCP4 (Purkinje cell protein 4). Cluster 3 included ITPR2 (inositol 1,4,5-trisphosphate receptor type 2), which is expressed in astrocytes and is involved in synapse elimination ^45^. GRIN2A, PCP4, and ITPR2 are also modulated in human ASD ^31^.

Gene expression alterations at 3M and 6M showed a positive correlation with those in human ASD ^31^. Contrastingly, gene expression alterations in marmosets at 3M were negatively correlated with those in iPSCs from patients with ASD. These gene expression modulations in the fetal period differ from those after birth ^32^. This suggests common temporal dynamics of abnormal gene expression between the model animal and human idiopathic ASD from the fetal and mature stages. However, gene expression modulations in marmosets at 0M were not correlated with those in mature patients or iPSC-derived neurons. This suggests that the perinatal period is a unique time point with a distinct pattern of gene expression modulations from other ages.

Synaptic alterations in VPA marmosets were inconsistent with previous studies on rodent VPA models. In VPA-exposed rodents, the spine density was lower from post-weaning to adult ages ^46^. Further, the E/I ratio was higher in cortical neurons ^47^, which is inconsistent with our results. Compared to VPA-exposed marmosets, the rat VPA model showed a weaker correlation of gene expression alterations with those in human ASD. The marmoset ASD model seems to reproduce human ASD more precisely than rodent VPA model, which supports the validity of this model for translational research.

There have been pharmacological interventions for patients with ASD after adolescence; however, they generally have short-term effects. Earlier intervention may modify the developmental trajectory and may achieve lifelong good outcome. Normalizing the expression of affected genes at appropriate time points could be an effective ASD therapy. GRIN2A expression upregulation ^48^ could ameliorate ASD symptoms by modulating downstream signals. Cholinesterase inhibitor donepezil has been applied in some patients with ASD ^49^ and could be more effective if administered soon after birth, because CHRM3 (cholinergic receptor muscarinic 3) was a cluster 1 gene and was only downregulated at the neonate stage. However, early-age intervention requires early screening and diagnosis; moreover, medication to neonates and children generally involves risks of adverse side effects.

The foundation for future learning, mental health, and life success are built in the early stage. Therefore, earlier intervention is desirable for developmental disorders, including ASD. This is the first report regarding altered biological characteristics (synapse and gene expression) in early-age ASD using non-human primates. Several rodent models have provided insights regarding ASD. However, non-human primates have a much closer dynamics as those for brain development with humans ^8,10,11^, and therefore could have greater potential in translational ASD research. VPA model marmosets have synaptic and molecular phenotypes close to human ASD, as well as autistic behavior and morphological alteration ^12,13,18,19^. There were distinct differences in the biological backgrounds at the neonate, childhood, and mature stages. The impaired synaptogenesis and circuit remodeling at the neonate and childhood stage could affect subsequent normal brain development since they are crucially involved in experience-dependent circuit formation. We did not address the roles of non-neuronal components such as astrocytes and microglia, of which accumulating evidence suggests the involvement in the pathogenesis of ASD. Nonetheless, this study indicates the critical importance of understanding early biological phenotypes of ASD. Further, its findings could contribute to future age-dependent therapy for ASD at early life, which could be highly efficient.

## Acknowledgments

We thank A. Tsuchiya for assisting the experiments. This work was supported by Intramural Research Grant for Neurological and Psychiatric Disorders from the National Center of Neurology and Psychiatry (29-6, NI), Novartis Research Grant (SW), JSPS KAKENHI Grant Number JP18K06497 (JN) and AMED grant number 19dm0207066h0001 (NI).

## Methods

### Animals

All experiments were approved by the Animal Research Committee of the National Center of Neurology and Psychiatry and the Animal Care and Use Committee of the National Institute of Radiological Sciences, and were in accordance with the NIH Guide for the Care and Use of Laboratory Animals. Marmosets were bred in the National Center of Neurology and Psychiatry or the National Institute of Radiological Sciences. Serum progesterone levels in female marmosets were measured; further, we determined the ovulation time. VPA was prepared as a 4% solution in 10% glucose solution and intragastrically administered to pregnant marmosets daily for 7 days from post-conception day 60 at 200 mg/kg/day. Some UE animals received 10% glucose solution, while other UE animals received no treatment. The offspring of VPA-administered or UE marmosets were used at 0 months (M) (4-11 days for electrophysiological and morphological analyses, 0-4 days for molecular analyses), 3M (88-110 days for electrophysiological and morphological analyses, 84-99 days for molecular analyses), and 6M (189-273 days for electrophysiological and morphological analyses, 180-208 days for molecular analyses). There were no between-group differences in the pregnancy periods of the glucose-administered UE and VPA marmosets (139.5 ± 1.2 days for glucose-administered UE [mean ± SD, n=6] and 140.6 ± 1.7 days for VPA [n=16], p=0.10 with *t*-test), as well as the body weight at birth (29.0 ± 1.9 g for glucose-administered UE and 30.7 ± 2.4 g for VPA, p=0.088). There was no significant difference in the body weight at birth between glucose-administered and untreated UE animals (29.7 ± 2.9 g for untreated animals, n = 36, p = 0.45).

### Slice preparation and electrophysiology

The marmosets were deeply anesthetized using ketamine hydrochloride (20 mg/kg, i.m.) and sodium pentobarbital (100 mg/kg, i.p.). Next, they were transcardially perfused with ice-cold CO_2_/O_2_-saturated artificial cerebrospinal fluid (ACSF); subsequently, the skull was removed and the brain isolated. The ACSF comprised of the following (in mM): 126 NaCl, 3 KCl, 1.2 NaH_2_PO_4_, 10 glucose, 26 NaHCO_3_, 2.4 CaCl_2_, and 1.3 MgSO_4_. Coronal slices containing the dorsomedial prefrontal cortex (area 8b/9) were prepared using a vibratome (model MT, Dosaka EM) at 400 µm thickness (Fig. 1a). Typically, 7-8 slices were obtained from each hemisphere. The slices were placed on an interface-style chamber perfused with ACSF at 32°C to allow recovery.

We obtained whole-cell recordings from layer 3 pyramidal neurons under an infrared differential image contrast (IR-DIC) microscope (Olympus, BX51WI). The slices were placed on a recording chamber that was continuously perfused with ACSF at 28°C. To record miniature synaptic currents, 1 µM tetrodotoxin (Fujifilm Wako Pure Chemical) was added to the ACSF. In some experiments, the AMPA receptor blocker NBQX (Abcam, 20 µM) or the GABA_A_ receptor blocker picrotoxin (Cayman Chemical, 100 µM) were added to the ACSF. The internal solution contained the following (in mM): 130 Cs methanesulfonate, 10 NaCl, 5 MgSO_4_, 10 HEPES, 0.6 EGTA, 2 Na-ATP, 0.6 Na-GTP, 10 Na-phosphocreatine, and 3 mg/ml biocytin. A patch-clamp amplifier (Axopatch 1D, Axon Instruments) was used for signal recording with a low-pass filter of 2 kHz. The signals were recorded on a PC using a data acquisition board (PCI-6221, National Instruments) and IgorPro software (WaveMetrics) at a 10 kHz sampling frequency. Data with a series resistance > 28 MΩ for 0M and > 18 MΩ for 3M and 6M were excluded. We recorded mEPSCs at −65 mV, which was close to the calculated equilibrium potential for Cl^-^, and mIPSCs at 0 mV, which was close to the equilibrium potential for monovalent cations. Miniature synaptic currents were analyzed using a custom-made program for MatLab (Mathworks) based on the algorithm used by Mini Analysis (Synaptosoft). The signal was further band-pass filtered at 4-1000 Hz. Moreover, the standard deviation (SD) of the baseline was measured in a 50-ms segment lacking events. We set the current amplitude threshold (in the units of SD of the baseline) and current area threshold (in the units of ms times SD of the baseline) for miniature events at 3 and 4.5 (0M mEPSCs), 3 and 9 (0M mIPSCs), 2 and 3 (3M and 6M, mEPSCs), 2 and 6 (3M and 6M, mIPSCs), respectively.

Stimulus-evoked synaptic currents were induced by field stimulation of layer 4-5 using a bipolar tungsten electrode (tip separation 150 µm) connected to an isolator (Dagan, BSI-950). The stimulus duration was 0.1 ms. The intensity was set to evoke about half-maximal EPSC (400 µA for 0M, 300 µA for 3M and 6M). EPSCs and IPSCs were recorded at a holding potential of -65 mV and 0 mV, respectively.

Stimulus-evoked field potentials were recorded using an interface-style recording chamber under a binocular microscope. The chamber was maintained at 35°C and a glass recording electrode containing 0.5 M NaCl was placed in layer 3 at about 400-500 µm from the pia. A bipolar tungsten stimulation electrode was placed in layers 4-5 at about 1-1.2 mm from the pia. The signal was amplified using an amplifier (Cygnus ER-1) with a low-pass filter at 1 kHz and recorded on a PC at a 10-kHz sampling frequency. Test stimuli were applied at a 30-s inter-stimulus interval. Up to two pathways were used in the same slice. The stimulus duration was 0.1 ms. The stimulus intensity was adjusted to evoke a field EPSP with one phase decay (100-500 µA). The field EPSP was completely abolished after perfusion with 20 µM NBQX (Tocris) and 50 µM D-APV (Cayman Chemical) (data not shown). When the baseline amplitude was stable for > 20 min, we applied LFS (1-s interval, 900 times) from the stimulation electrode. The dependency of LFS-induced LTD on NMDA receptors was confirmed using 100 µM D-APV in 0M slices. We induced mGluR-dependent LTD by perfusion with 100 µM DHPG (Tocris) for 10 min.

### Neuron structure analysis

After whole-cell recording using biocytin-containing electrodes, the slices were fixed in 4% paraformaldehyde overnight and stained using Vectastain Elite ABC Kit (Vector Laboratories) and DAB Substrate Kit (Vector Laboratories). The slices were dehydrated using a graded ethanol series, cleared with xylene, and mounted with Entellan New (Merck).

The neuron structure was analyzed using Neurolucida (MBF Bioscience). The whole dendritic arbor was traced using a 20 × objective. The dendritic spine density was analyzed using a 100 × oil immersion objective (NA 1.49) on the basal dendrites at 25-50 µm from the soma for 0M and at 50-75 µm from the soma for 3M and 6M, which corresponded to the highest spine density ^10^. We used an intensity-based method to calculate the spine volume given that direct measurement of the spine diameter is unreliable for small spines. Axial and transversal profiles of 3-6 medium-sized spine heads on a dendrite were fitted using a Gaussian curve (Suppl. Fig. 1). For spines with a diameter > 0.4 µm, twice the standard deviation of the curve (2σ) matches the actual spine head diameter observed using ultra-high voltage electron micrograph (Oga, T. and Fujita, I., unpublished observation). Therefore, we used 2σ as the diameter. The spine volume was calculated using the transversal (*d*_t_) and axial diameter (*d*_a_) as π*d*_t_^2^*d*_a_/6. We obtained a proportional relationship between the calculated spine volume and optical density for spine heads (diameter > 0.4 µm and volume < 0.4 µm^3^) as references (Extended Data Fig. 5). Using this relationship, the other spine volumes were estimated by their optical density measurements.

For electron microscopy, slices were immediately fixed after slicing in 2.5% glutaraldehyde and 2% paraformaldehyde in 0.1 M cacodylate buffer (pH 7.2) for at least 5 days. The slices were then treated using 1% osmium oxide in cacodylate buffer for 60 min and stained with 2% uranyl acetate for 60 min. After serial dehydration, the tissues were embedded in Epon812 (TAAB). Ultra-thin sections (70 nm thick) were prepared and examined using a Tecnai Spirit electron microscope (Thermo Fisher Scientific-FEI).

### Analysis of vocalization

Marmoset vocalizations were recorded using a linear PCM recorder (Olympus, LS-100). The marmosets were isolated in a soundproof chamber and were allowed habituation to the environment for approximately 1 min before each recording session. The animals were placed in a small recording cage (38 × 43 × 47 cm^3^) located 10 cm apart from the recorder. Further, vocalizations uttered during 5-min recording sessions were obtained in 24 bit at a 96-kHz sampling frequency. The recordings were started from postnatal 1-3 weeks and were performed every week up to 20 weeks, of which the data from 9-11 weeks were analyzed. The data were analyzed by an experimenter who did not know the type of the animal, using the Praat software ^50^. Each call was annotated as previously described ^51–53^. Continuous sequences of multiple calls without a silent gap (multisyllabic calls) were considered as a single bout of call. Call type entropy was calculated using the ratio of each call type *r*_*i*_ as −Σ_*i*_ *r*_*i*_ log_2_ *r*_*i*_.

### Microarray analysis

Microarray analysis was performed as previously described ^54,55^. In addition to the two prefrontal areas (areas 8 and 12), we used temporal association area (area TE) to increase reliability of data. Marmosets at 0M, 3M, and 6M were anesthetized using ketamine hydrochloride and sodium pentobarbital. Subsequently, they were transcardially perfused with diethylpyrocarbonate-treated phosphate-buffered saline followed by isolation of the cortical tissues and immersion in RNAlater (Thermo Fisher). Total RNA was extracted using the RNeasy Mini Kit (Qiagen). RNA integrity was assessed using the Bioanalyzer (Agilent Technologies) and samples with a RIN value > 7 were used. A biotin-labeled cRNA probe was generated using GeneChip 3’IVT Express Kit (Affymetrix). The probe was hybridized with a custom-made microarray (Marmo2a520631F) ^54,55^ using the GeneChip Hybridization, Wash, and Stain Kit (Affymetrix). The microarray was scanned using the GeneChip Scanner 3000 (Affymetrix), and processed using MAS5 (Affymetrix) to examine the reliability of probe detection. The data were normalized using the GCRMA software (Bioconductor). We considered genes with log_2_ expression value > 5 as being expressed in the brain tissue. Protein-coding genes were selected based on the HUGO Gene database (https://www.genenames.org). Differential gene expression was evaluated based on the p-value from Welch’s *t*-test with Benjamini-Hochberg adjustment (p_adj_). DEGs (log_2_ expression value > 6, absolute value of logFC > 0.4 and p_adj_ < 0.05 at either age) were clustered using the *k*-means algorithm. The cluster number was set to 3, which yielded the lowest value of Akaike’s information criterion. For affected genes with multiple probes, we used data from the probe with the lowest p_adj_.

The mean distance between the logFC of synaptic phenotypes (logFC_syn_) and gene expression (logFC_gene_) was calculated as Σ_*t*_ | logFC_syn_(*t*) − logFC_gene_(*t*)| / 3, where *t* is the age (0M, 3M, or 6M). Pathway analysis was conducted using the IPA software (Qiagen). The list of ASD-related genes was derived from the Simons Foundation Autism Research Initiative (SFARI, http://www.sfari.org, release 08-07-2020). The list of critical period-related genes was obtained by combining data from two studies ^29,30^. To compare gene expression modulations with those in human ASD (postmortem samples^31^ or iPSC-derived neurons ^33^), and commonly modulated genes with p_adj_ < 0.1 were used. To compare gene expression modulations in rat ASD model ^34^ with those in human ASD, gene symbols were converted using BioMart (www.ensembl.org), and commonly modulated genes genes with p_adj_ < 0.05 were used.

### Statistical analyses

To compare UE and VPA data at multiple time points, the data were tested using the Kolmogorov-Smirnov test for normality and the Levene test for variance homogeneity. To satisfy the requirements for variance homogeneity, the frequencies and amplitudes of miniature synaptic currents were transformed to the power of 0.2, and the E/I ratio of miniature and evoked (0M only) synaptic currents were transformed to the inverse. Subsequently, the data were tested with two-way ANOVA, followed by the Holm-Sidak test. Two-way repeated measures ANOVA was used to compare the paired-pulse ratio. Between-group comparisons were performed using the *t*-test. The Kolmogorov-Smirnov test was used to compare the distribution of spine volume and PSD length. Enrichment of SFARI genes, critical period-related genes, and genes belonging to human ASD-related modules in each gene cluster were analyzed using Fisher’s exact test. Spearman’s correlation was used to determine the correlation of logFC values between marmosets or rats and humans. All the data were from distinct samples. Tests for gene enrichment were one-sided, and all other tests were two-sided. Error bars in the graphs represent SEM.

---

**Supplementary Table 1**. List of affected genes categorized into three clusters. The logFC and p_adj_ values for each time point, as well as the module, logFC, and p_adj_ in human ASD ^31^, are shown.

**Supplementary Table 2**. Critical period-related genes listed in two studies ^29,30^ and expressed in the marmoset brain. For marmoset DEGs, cluster number is shown.

**Supplementary Table 3**. Pathway analysis of the gene clusters.

**Extended Data Fig. 1.**
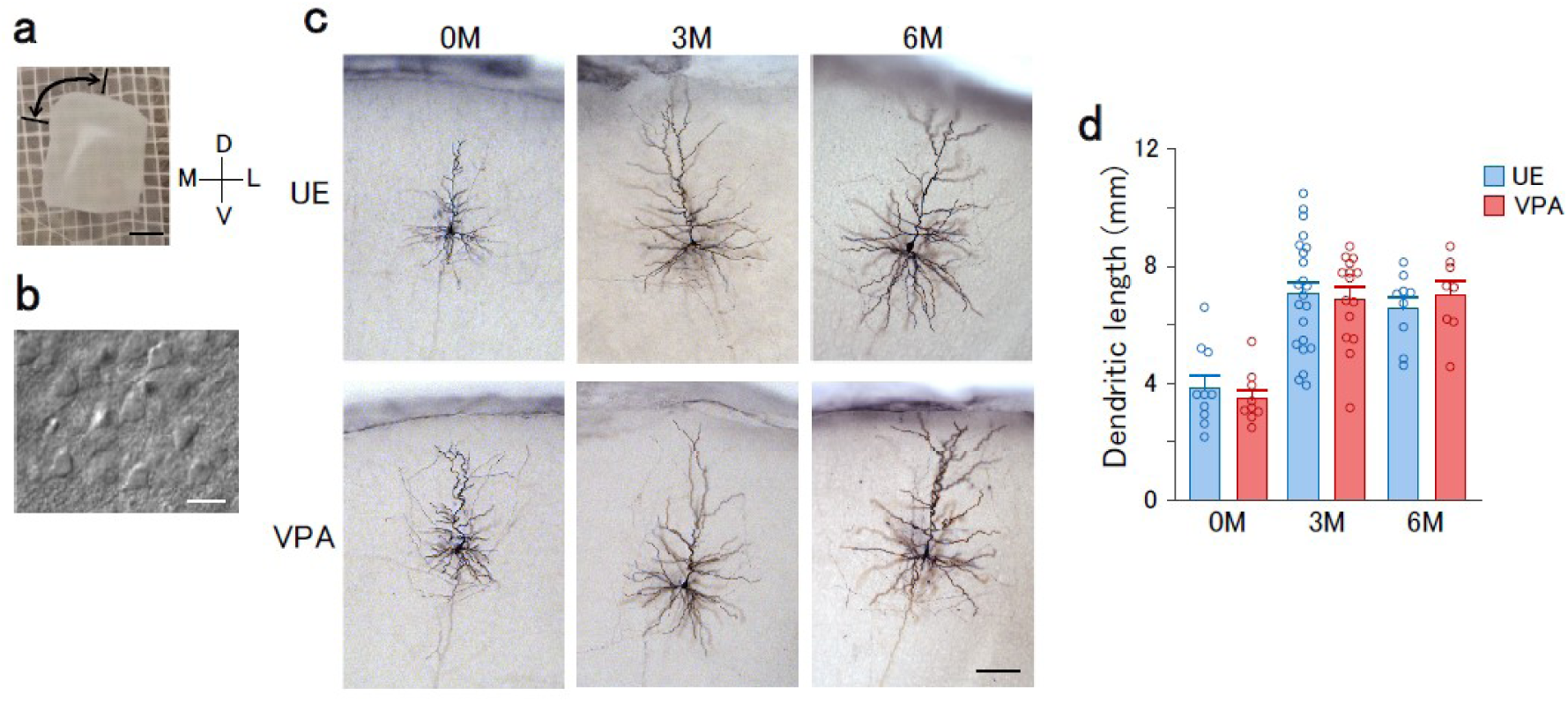
Acute slice preparation and development of dendrites in layer 3 pyramidal neurons in the dorsomedial prefrontal cortex. **a**. Representative slice preparation of the prefrontal cortex of a marmoset. The dorsomedial part (arrow) corresponds to area 8b. Scale bar, 2 mm. M, medial; L, lateral; D, dorsal; and V, ventral. **b**. IR-DIC image of layer 3 of the slice, which shows the pyramidal neuron somata. Scale bar, 20 µm. **c**. Photomicrographs of biocytin-stained layer 3 pyramidal neurons from UE and VPA animals at 0M, 3M, and 6M. Scale bar, 100 µm. **d**. Total dendritic length of layer 3 pyramidal neurons from UE (blue) and VPA (red) animals at 0M, 3M, and 6M. Main effect of treatment, F(1,67) = 0.0025, p = 0.96; interaction of age × treatment, F(2,67) = 0.34, p = 0.71. Number of data: 10 cells in 4 animals (0M UE), 9 cells in 2 animals (0M VPA), 22 cells in 4 animals (3M UE), 15 cells in 3 animals (3M VPA), 9 cells in 3 animals (6M UE), and 8 cells in 2 animals (6M VPA).

**Extended Data Fig. 2.**
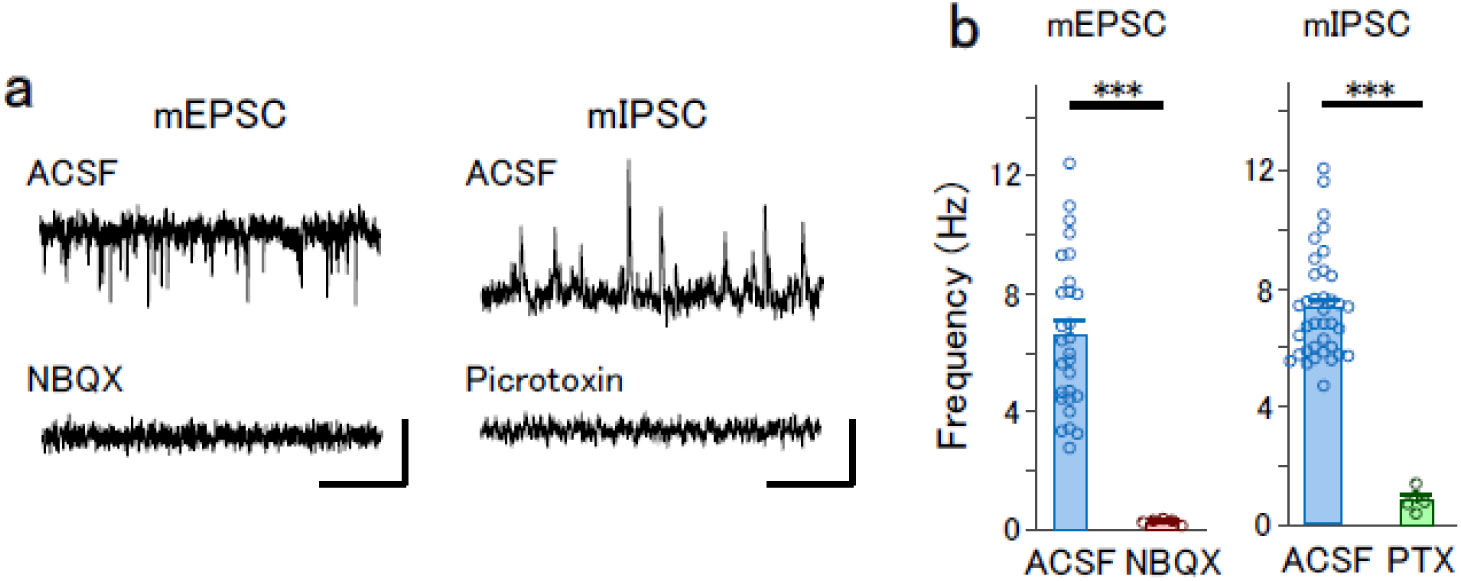
Effects of AMPA and GABA_A_ receptor blockers on miniature synaptic currents in UE animals at 3M. **a**. Representative traces of mEPSCs in normal ACSF and in the presence of NBQX (left), as well as mIPSCs in normal ACSF and in the presence of picrotoxin (right). **b**. NBQX effects on the frequency of mEPSCs (left; *t*-test, ***p = 6.9 × 10^−11^) and picrotoxin effects on the frequency of mIPSCs (right; ***p = 1.6 × 10^−22^). Number of data: 5 cells in 2 animals (mEPSCs, NBQX) and 5 cells in 2 animals (mIPSCs, picrotoxin). The data in ACSF are the same as in Fig. 1 (mEPSC) and Fig. 2 (mIPSC).

**Extended Data Fig. 3.**
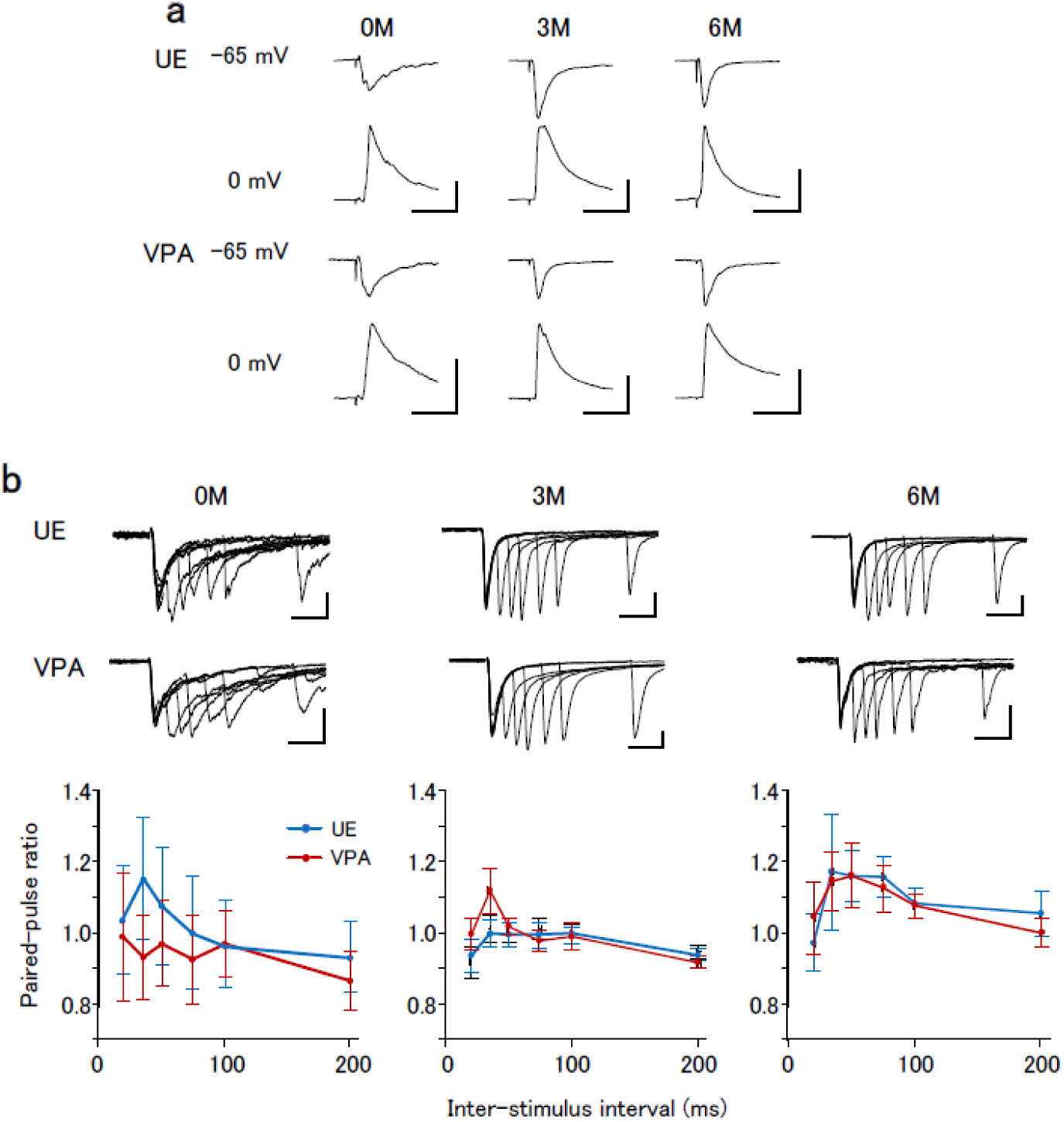
Evoked EPSCs and IPSCs by stimulation of layers 4-5, as well as effects of VPA exposure on the paired-pulse ratio of evoked EPSCs. **a**. Representative traces of stimulus-evoked EPSCs and IPSCs in UE and VPA animals at 0M, 3M, and 6M. Stimulus intensity: 400 μA for 0M and 300 μA for 3M and 6M. Vertical scale bars: 200 pA for EPSCs and 400 pA for IPSCs at 0M; 500 pA for EPSCs and 1000 pA for IPSCs at 3M and 6M. Horizontal scale bars, 50 ms. **b**. EPSC evoked by paired-pulse stimulation (inter-stimulus interval 20-200 ms) in UE (top) and VPA (bottom) animals at 0M, 3M, and 6M. Vertical scale bars: 50 pA for 0M; 200 pA for 3 and 6M. Horizontal scale bar, 50 ms. The paired-pulse ratio in UE (blue) and VPA (red) animals are plotted as a function of the inter-stimulus interval. Repeated measures ANOVA, main effect of treatment for 0M, 3M, and 6M, F(1,25) = 0.17 (p = 0.69), F(1,35) = 0.69 (p = 0.41), and F(1,21) = 0.0002 (p = 0.99), respectively; interaction of intensity × treatment, F(4,100) = 0.77 (p = 0.55), F(4,140) = 1.96 (p = 0.10), and F(4,84) = 0.30 (p = 0.88), respectively. Number of data: 17 cells in 6 animals (0M UE), 10 cells in 2 animals (0M VPA), 23 cells in 5 animals (3M UE), 14 cells in 2 animals (3M VPA), 12 cells in 2 animals (6M UE), and 11 cells in 2 animals (6M VPA).

**Extended Data Fig. 4.**
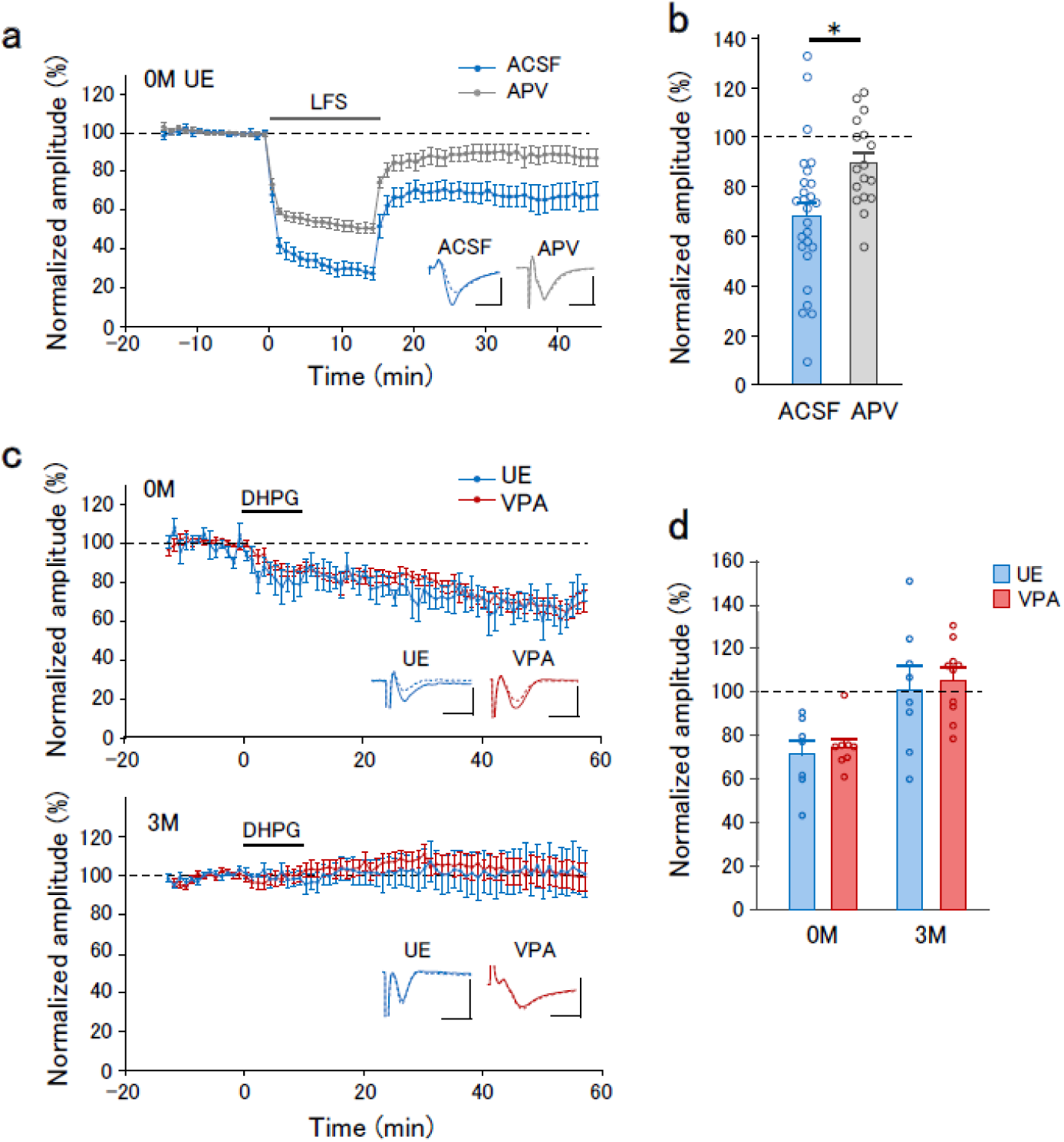
Dependency of LFS-induced LTD on APV and effects of VPA exposure on mGluR-dependent LTD. **a**. The time course of the normalized field EPSP amplitudes in UE animals at 0M in normal ACSF (blue) and in APV presence (gray). Insets represent the average traces before (−10 to 0 min, solid lines) and after (30 to 40 min, dotted lines) LFS. **b**. Normalized EPSP amplitudes in normal ACSF and APV presence after LTD induction (30 to 40 min). *t*-test, *p = 0.030. Number of data: 26 pathways in 22 slices from 12 animals (ACSF); 17 pathways in 13 slices from 3 animals (APV). **c**. The time course of the normalized EPSP amplitude in UE (blue) and VPA (red) animals at 0M. LTD was induced by perfusion of DHPG (100 µM) for 10 min. Insets represent the average traces before (−10 to 0 min, solid lines) and after (30 to 40 min, dotted lines) DHPG application. **d**. Normalized EPSP amplitudes at 30-40 min after the onset of DHPG perfusion. Two-way ANOVA: main effect of treatment, F(1,29) = 0.27, p = 0.61; interaction of age × treatment, F(1,29) = 0.002, p = 0.97. *t*-test between 0M and 3M in UE animals, p = 0.029. Number of data: 7 pathways in 4 slices from 2 animals (0M UE), 8 pathways in 5 slices from 1 animal (0M VPA), 8 pathways in 5 slices from 3 animals (3M UE), and 10 pathways in 5 slices from 3 animals (3M VPA).

**Extended Data Fig. 5.**
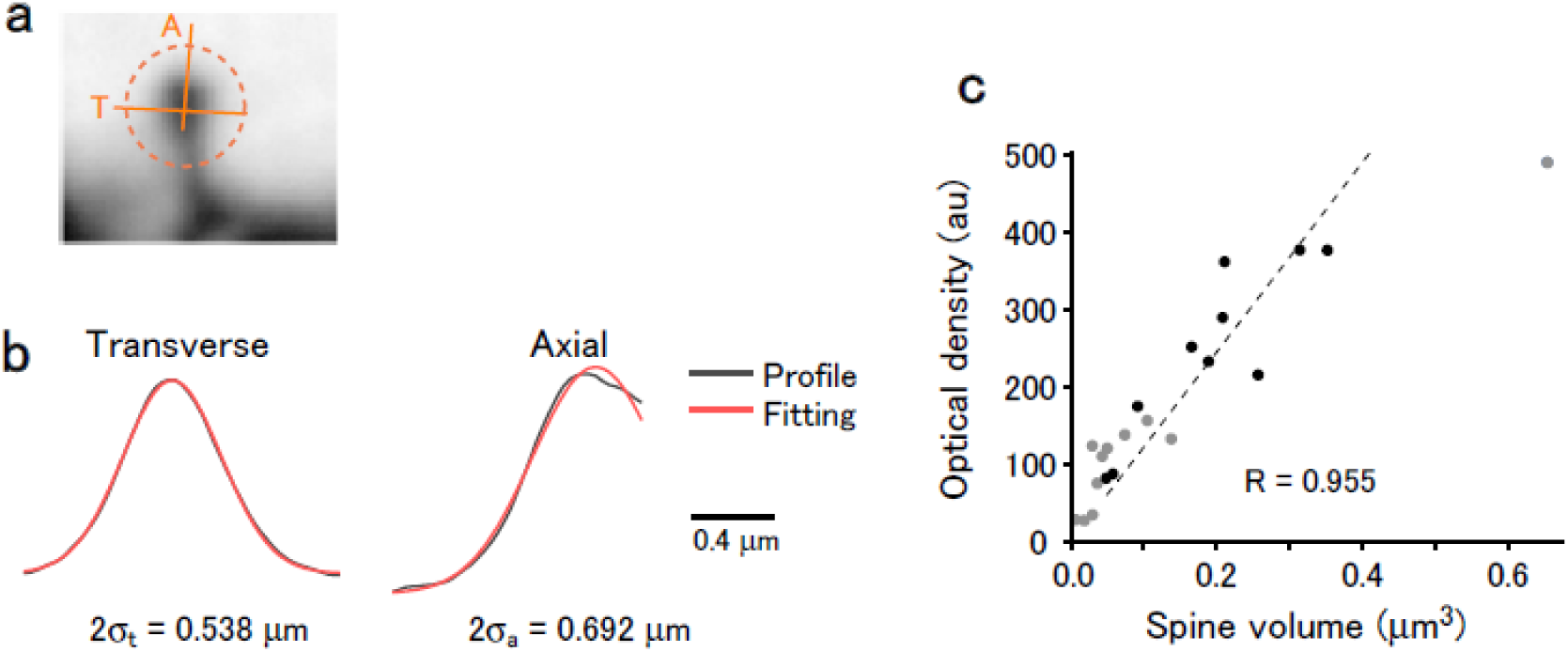
Measurement of spine volumes in biocytin-filled neurons. **a**. Representative image of a dendritic spine. Optical density profiles along the transverse (T) and axial (A) lines are shown in (b). Scale bar, 1 µm. **b**. Optical density profiles along the lines shown in (a) (black) and their Gaussian fitting (red). **c**. Relationship between the calculated spine volume and the optical density in 20 spines on a dendrite. The dotted line represents the linear regression between the optical density and volume for spines with a diameter > 0.4 µm and volume < 0.4 µm^3^ (black dots).

## Notes

### Competing Interest Statement

The authors have declared no competing interest.

